# Probabilistic neural code supports metacognition in value-based decisions in the human brain

**DOI:** 10.64898/2026.04.21.719717

**Authors:** Ling Huang, Hui-Kuan Chung, Hsin-Hung Li

**Affiliations:** Department of Psychology, The Ohio State University, Columbus, OH 43201, USA; Department of Economics, University of Zurich; Zurich, 8006, Switzerland; Zurich Center for Neuroeconomics, University of Zurich; Zurich, 8006, Switzerland

## Abstract

Subjective value underlies many everyday decisions, yet the neural representations supporting these valuations are inherently noisy and inevitably subject to uncertainty. Whether humans have metacognitive access to this uncertainty, and how such access is supported neurally, remains unclear. Here, we provide converging behavioral and neural evidence that subjective value is represented probabilistically in the human brain. Behaviorally, participants’ explicit uncertainty reports tracked trial-by-trial fluctuations in valuation variability. Neurally, we decoded full probability distributions over value from fMRI population activity in ventromedial prefrontal cortex (vmPFC) and superior parietal lobe and intraparietal sulcus (SPL-IPS). Decoded neural uncertainty predicted behavioral reported uncertainty at the trial, item, and individual levels. These findings indicate that neural populations jointly encode value and uncertainty, extending probabilistic population coding from perception and working memory to subjective value, and providing a neural basis for metacognitive access to uncertainty in value-based decisions.

## Introduction

Value-based decision-making requires the brain to assign subjective values to options and use these values to guide behavior (Bartra et al., 2013; Chib et al., 2009; Chung et al., 2025; Grabenhorst & Rolls, 2011; Harris et al., 2011; Kahnt, 2018; Padoa-Schioppa & Assad, 2006; Plassmann et al., 2007; Rich & Wallis, 2016; Rushworth et al., 2011). The cognitive and neural processes underlying subjective value are inherently noisy and variable. Even for deterministic and familiar items, an individual’s valuation fluctuates across repeated encounters. Decisions based on such noisy representations are therefore inevitably uncertain and may reflect imprecise estimates of one’s true preference (Barretto-García et al., 2023; Frydman & Jin, 2022; Kable & Glimcher, 2009; Khaw et al., 2021; Padoa-Schioppa, 2013; Polanía et al., 2019; Rangel & Hare, 2010; Woodford, 2020; Zylberberg et al., 2024). Access to this uncertainty, a form of metacognition, allows individuals to know how much they can trust their own valuation, and to guide actions, such as choosing the option with the highest value, seeking additional information or deferring a decision when the internal estimates are too unreliable (De Martino et al., 2013; Fleming, 2024; Lee & Daunizeau, 2021; Pouget et al., 2016; Walker et al., 2023).

Despite this, it remains unclear how value uncertainty is represented in the brain and whether individuals have metacognitive access to it. Prior studies of metacognition in value-based decisions have predominantly focused on confidence in discrete choice tasks, showing that confidence report tracks choice accuracy or stochasticity, although confidence itself can arise from multiple sources (Brus et al., 2021; da Silva Castanheira et al., 2021; De Martino et al., 2013; Folke et al., 2016; Landron et al., 2025; Plate et al., 2025). However, the extent to which individuals can similarly evaluate the uncertainty in their continuous value estimates remains underexplored , despite its importance for real-world decisions that require continuous valuation, such as deciding how much to invest. Importantly, investigating metacognitive access to continuous valuation speaks to a more fundamental question regarding the nature of value representation. If individuals can track trial-by-trial fluctuations in the quality of valuation, this would suggest that value is not represented merely as a point estimate, but as a richer representation that provides uncertainty for guiding adaptive behavior.

A normative framework of the neural basis for such a probabilistic representation was initially developed for perceptual and sensorimotor domains. The theory of probabilistic population codes posits that neural populations encode a probability distribution over the stimulus space, by incorporating the knowledge of the signal and noise structure of the population neural response (Foldiak et al., 1993; Jazayeri & Movshon, 2006; Ma et al., 2006; Sanger, 1996; Zemel et al., 1998). Such a neural code provides a joint representation of the estimate and uncertainty of a stimulus, and it has been proposed to underlie a range of Bayesian-like behaviors, including decision-making, planning, and multisensory or cue integration, by enabling the brain to weight information according to its reliability (Beck et al., 2008; Knill & Pouget, 2004; Ma et al., 2006; Walker et al., 2020). Moreover, it provides a potential neural basis for metacognitive judgments, allowing individuals to explicitly report the uncertainty of their own estimates. In support of this framework, empirical studies have shown that probabilistic information decoded from early visual cortex can predict the uncertainty of visual perception (Chetverikov & Jehee, 2023; Van Bergen et al., 2015; Walker et al., 2020). Recently, we demonstrated a similar neural coding scheme for higher-level cognitive processes such as visual working memory in human cortex (Li et al., 2021, 2025). Whether the same computational principles underlie the neural representation of subjective value in the human brain remains an open question (see a recent study investigating relevant research questions in vmPFC (Bouc et al., 2026)).

In this study, we first adapted a behavioral task where participants explicitly reported uncertainty for each food item after providing their subjective value estimate (Becker et al., 1964; Dost & Wilken, 2012; Wang et al., 2007). By testing the same item with multiple repetitions, we demonstrated behaviorally that reported uncertainty tracks trial-by-trial error (or variability) in continuous valuation, establishing that people have metacognitive access to their value estimates. Next, we measured population neural response by functional magnetic resonance imaging (fMRI). We focused on the brain regions that have been indicated in representing value and involved in value-based decision-making, including ventromedial prefrontal cortex (vmPFC; Bartra et al., 2013; Chung et al., 2025; Grabenhorst & Rolls, 2011; Harris et al., 2011; Kahnt, 2018; Padoa-Schioppa & Assad, 2006; Rich & Wallis, 2016; Rushworth et al., 2011), parietal cortex (superior parietal lobe and intraparietal sulcus; SPL-IPS; Chung et al., 2017, 2025; Grabenhorst & Rolls, 2011; Harris et al., 2011; Kahnt, 2018; Platt & Glimcher, 1999; Rushworth et al., 2011) and ventral striatum (Bartra et al., 2013; Chung et al., 2017, 2025; Harris et al., 2011; Kahnt, 2018; Knutson et al., 2001). By building an encoding- and-decoding model for BOLD activity (Van Bergen et al., 2015; Van Bergen & Jehee, 2021), we tested the hypothesis that probability distribution over subjective value can be decoded from value-related brain regions, and the decoded distributions predict both valuation and behavioral report of uncertainty.

## Results

### Humans track trial-by-trial value uncertainty

In Experiment 1, to examine whether participants had metacognitive access to trial-by-trial uncertainty in their valuations, we adapted an incentive-compatible Becker–DeGroot–Marschak (BDM) procedure (Becker et al., 1964) in which participants reported both a willingness-to-pay (WTP) estimate and an uncertainty range for different food items (common salty and sweet snack foods). Specifically, participants reported uncertainty by setting a maximum and a minimum price surrounding the reported WTP on the price scale, and the range was defined as the distance between the two (Figure 1a). Similar procedures have been used to measure uncertainty in value (Dost & Wilken, 2012; Wang et al., 2007) and visual memory (Honig et al., 2020; Li et al., 2021; Yoo et al., 2018).

**Figure 1.**
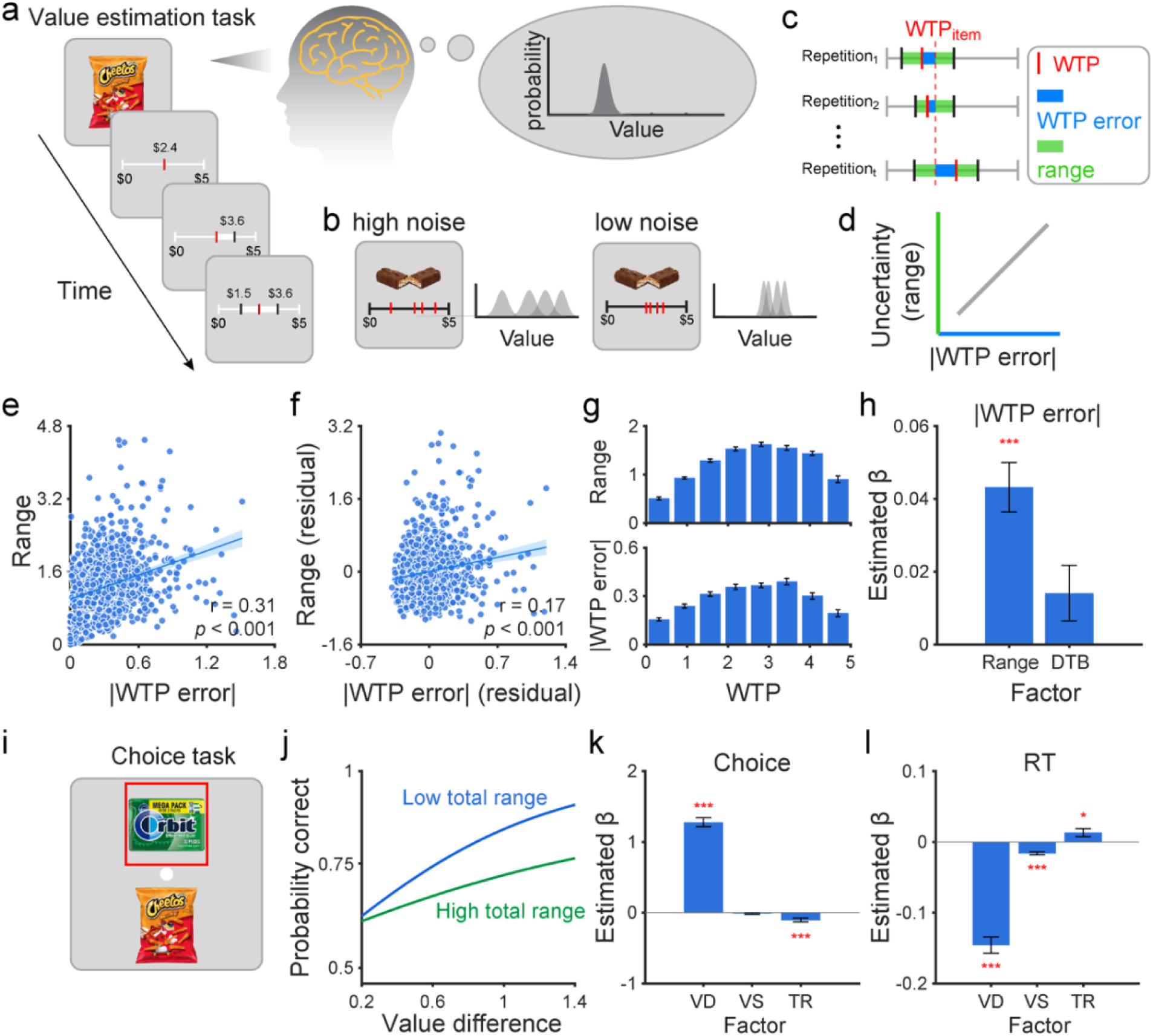
Procedures and behavioral results. (a) Subjective value estimation task procedure. Left: Each trial began with a 6 s food image presentation, followed by a 10 s response window during which participants reported their willingness to pay (WTP) and an uncertainty range on a continuous $0–$5 scale by setting a maximum and a minimum bound. (see Methods). Right: Schematic of the probabilistic population code hypothesis. Neural populations represent the subjective value of an item as a probability distribution over the value space, allowing a joint representation of the value estimate and the associated uncertainty. Readout of the mean of the probability distribution guides WTP reports, and the spread of the probability distribution reflects one’s valuation uncertainty, as captured by the reported uncertainty range. (b) Probabilistic representations track the fluctuations of the quality of valuation (even for the same item). High internal noise leads to representations with higher uncertainty (wide distributions) and variable estimates (WTPs indicated by the red marks). Low internal noise leads to more precise representations and less variable estimates. (c) Key behavioral measures. Item-level WTP (WTPitem) was defined as the mean WTP across repetitions and served as each item’s true value; trial-level WTP error as the absolute deviation of each trial’s WTP from WTPitem; and the trial-level uncertainty range was defined as the interval between the minimum bound and the maximum bound set by the participants. (d) Probabilistic representations allow participants to have metacognitive access to their valuation uncertainty, indicated by a positive correlation between WTP error and range. (e) A significant positive correlation between WTP error and reported Range pooled across all trials and participants (r = 0.31, p < 0.001). (f) Partial correlation between WTP error and reported Range after regressing out boundary distance from both variables (r = 0.17, p < 0.001), confirming that the relationship is not solely driven by boundary-induced compression near the scale endpoints. (g) Mean reported Range (top) and WTP error (bottom) as a function of WTP bin, each exhibiting an inverted-U relationship with item value. (h) Estimated fixed-effect coefficients (β ± SEM) from the linear mixed-effects model (LMM) predicting WTP error from reported range and distance to boundary (DTB), controlling for random intercepts of participants, items, and their interaction. Reported range significantly predicted WTP error (β = 0.043 ± 0.007, p < 0.001), demonstrating that the relationship between range and WTP error cannot be attributed to boundary proximity alone. *** p < 0.001. Error bars denote ±1 SEM across participants. (i) Example trial from the 2AFC choice task, in which two food items were presented simultaneously in the upper and lower visual fields and participants selected their preferred item, indicated by a red frame around the chosen item. (j) Illustration of the predicted effect of total range (TR) on choice behavior. Higher valuation uncertainty (high TR, green) is associated with a shallower psychometric curve, reflecting a reduced probability of choosing the higher-valued item across value differences, compared to low TR (blue). (k) Estimated fixed-effect coefficients (β ± SEM) from the generalized linear mixed-effects model (GLMM) predicting choice accuracy from value difference (VD), value sum (VS), and total range (TR). VD positively predicted choice accuracy, whereas TR negatively predicted accuracy independently of VD and VS. (l) Estimated fixed-effect coefficients (β ± SEM) from the LMM predicting response time (RT) in choice task from VD, VS, and TR. Larger VD and higher VS were associated with faster choices, whereas higher TR predicted longer RT. *p < 0.05, *** p < 0.001. Error bars denote ±1 SEM across participants.

Unlike domains with an objective ground truth, where metacognitive ability can be quantified by relating uncertainty or confidence reports to the magnitude of estimation error, subjective valuation lacks an explicit reference. Here, we hypothesized that meaningful uncertainty reports would track internal noise in value signals, the fluctuations in one’s value estimate across repeated evaluations, even for the same item (Figure 1b). To test this, participants evaluated each item four times (missing rates: WTP: 0.02 ± 0.02%; uncertainty report: 0.34 ± 0.11%; Supplementary Figure 1), allowing the variability of valuation and uncertainty report to be related. For each food item, the mean across the repetitions represents each participant’s stable preference (Figure 1c). Trial-level variability is then defined as WTP error, computed as the absolute difference between a single-trial WTP report and the corresponding item-wise mean, reflecting momentary fluctuations around each participant’s stable preference for that food item. With a probabilistic representation of value, metacognitive access to one’s own valuation would be reflected in higher reported uncertainty tracking the magnitude of these trial-by-trial WTP errors (Figure 1b&d).

Pooling across all trials and participants we observed a significant positive correlation between reported uncertainty (range) and the magnitude of WTP error (r = 0.31, p < 0.001; Figure 1e). One contributor to this relationship is the boundary of the rating scale. Both WTP error and reported range exhibited an inverted-U pattern as a function of WTP (Figure 1g), consistent with previous studies (Bobadilla-Suarez et al., 2020; Clairis & Pessiglione, 2022; Lebreton et al., 2015; Lopez-Persem et al., 2020; Plate et al., 2025; Polanía et al., 2019). This pattern likely arises from two factors. First, both WTP and range reports are constrained by the bounds of the rating scale, such that values cannot exceed these limits. Second, if value estimates are represented probabilistically, the corresponding belief distributions may be affected or truncated at the boundaries, which can lead to higher certainty at extreme values (Hahn & Wei, 2024; Lebreton et al., 2015).

To investigate whether uncertainty reports track WTP error beyond the effect of the boundary, we defined the boundary distance as the absolute distance between the WTP and the nearest scale boundary ($0 or $5). We found that the correlation between range and WTP error remained significant even when we first regressed out the boundary distance from both variables (r = 0.17, p < 0.001; Figure 1f). To target the uncertainty and variability within the same item, we fitted a linear mixed-effects model (LMM) to predict the magnitude of WTP error from reported range, controlling for boundary distance, the identity of the participant and the item (including an interaction term between item and participant). Reported range significantly predicted WTP error (β = 0.043 ± 0.007, p < 0.001; Figure 1h), demonstrating that participants had metacognitive access to trial-by-trial valuation variability beyond the effects of boundary and item-specific uncertainty.

Having established that participants had metacognitive access to their valuation uncertainty, we next asked whether the reported item-level uncertainty in the estimation task predicted choice behaviors in a different task. We asked the same participants to perform a 2AFC choice task (Figure 1i). Participants chose their preferred item between two items sampled from those tested in the estimation task (see Methods). To quantify uncertainty associated with each choice pair, we calculated total range (TR) as the sum of the two items’ mean reported range in the estimation task. Choice accuracy was defined as selecting the item with the higher item-wise

WTP based on the estimation task. TR significantly and negatively predicted choice accuracy (β = −0.104 ± 0.028, p < 0.001; Figure 1j&k) beyond value difference (VD; β = 1.277 ± 0.062, p < 0.001) and value sum (VS; β = −0.012 ± 0.010, p = 0.244), indicating that pairs with higher uncertainty were harder to discriminate. As a sanity check, replicating a previous study (Polanía et al., 2019), higher total WTP variability (TV), defined as the sum of the standard deviations of WTP reports across repeated presentations of the same item, similarly predicted lower choice accuracy (β = −0.459 ± 0.044, p < 0.001; Supplementary Figure 2).

Furthermore, to examine whether valuation uncertainty is reflected in choice decisiveness, we tested whether reported range in the estimation task predicted the response time in the subsequent choice task. We found that higher TR was associated with longer response time (β = 0.013 ± 0.006, p = 0.017; Figure 1l). Together, these findings demonstrate that item-level subjective uncertainty reports from the value estimation task captured meaningful valuation signals that predicted both subsequent choice and decisiveness.

### Subjective value can be decoded from vmPFC and parietal cortex

In Experiment 2, participants performed the same value estimation task of Experiment 1 while undergoing fMRI scanning. Here, each item was repeated 10-12 times (trials) in total across two sessions for each participant (see Methods). Behavioral results from the fMRI session showed high compliance (missing rates: WTP: 0.04 ± 0.03%; Range: 0.98 ± 0.29%; Supplementary Figure 1) and closely replicated those observed in Experiment 1. Reported uncertainty range significantly correlated with the magnitude of WTP error, even after the effect of the scale boundary and the identity of the items were controlled for (Supplementary Figure 3).

Given that participants can meaningfully report uncertainty in their subjective valuation trial-by-trial, here we test whether neural populations in value-related brain regions encode probabilistic representations of value that can be decoded from fMRI BOLD signals and jointly predict value and uncertainty report at a single-trial level. Single-trial BOLD response for valuation was obtained by modeling each 6 s food item presentation as a regressor for each trial. Critically, behavioral response timing was modeled as a separate regressor, ensuring that the signals we analyzed only reflect valuation rather than motor or response signals. In addition, in this experiment, the direction of the rating scale was flipped between the two scanning sessions for each participant, so the decoded value information cannot be attributed to spatial visual information or specific visual landmarks related to the rating scale.

We decoded probabilistic information of value from BOLD response by adapting a generative model-based decoding framework (Van Bergen et al., 2015; Van Bergen & Jehee, 2021). In a cross-validation procedure, in the training phase, we first fit a generative (encoding) that mapped subjective value (WTP) to multivoxel activity patterns modeled as a multivariate normal distribution (Figure 2a). The mean of the distribution was determined by each voxel’s value response function, approximated by a weighted sum of eight Gaussian basis functions evenly tiling the value space ($0–$5). The noise covariance was estimated by an empirical covariance matrix with shrinkage toward a theoretical covariance structure (see Methods). In the testing phase, we inverted this model by Bayesian inference to decode value from voxel activity on each trial, yielding a posterior probability distribution over the value space (Figure 2a). The mean of this posterior served as the decoded value estimate (decoded WTP), and its entropy as the decoded uncertainty (Chetverikov & Jehee, 2023).

**Figure 2.**
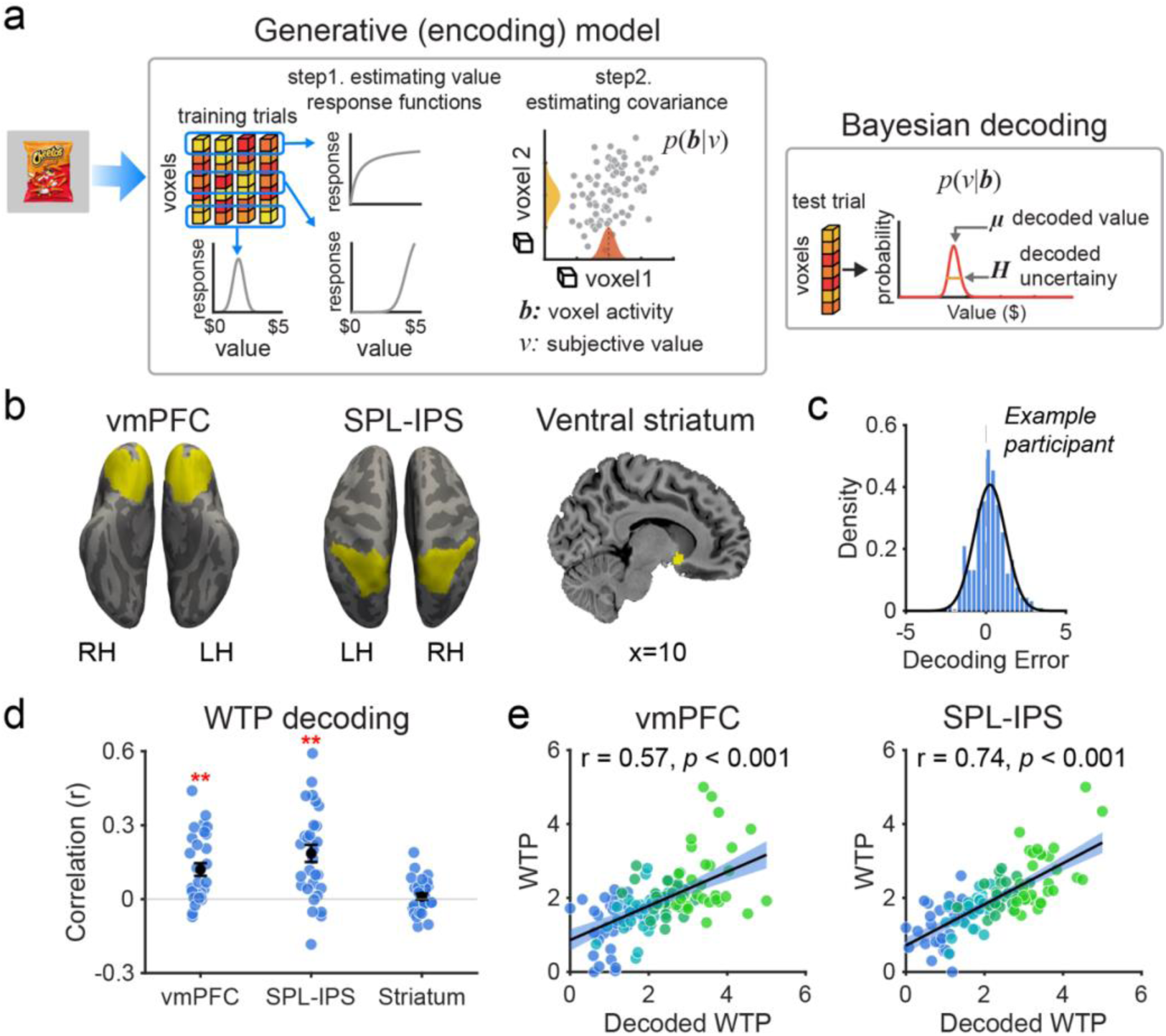
Subjective value can be decoded by a generative model-based decoding approach. (a) Schematic of the generative model-based decoding framework. Generative (encoding) model: Using training trials, a value response function was estimated for each voxel (step 1); next, the noise covariance structure of voxel activity was estimated (step 2), yielding a probability distribution *p*(**b**|*v*) over voxel activity pattern **b** given subjective value *v*. Bayesian decoding: For each test trial, based on the voxel activity pattern, a posterior probability distribution over value space, *p*(*v*|**b**) was decoded by inverting the generative model according Bayes’ rule. The mean of this posterior (*μ*) served as the decoded value estimate, and the entropy (H) of the posterior served as the decoded uncertainty, reflecting the spread of the distribution. (b) Three regions of interest (ROIs) were defined: vmPFC, SPL-IPS, and ventral striatum (highlighted in yellow). (c) Decoding error distribution in vmPFC for an example participant. (d) Pearson correlation between decoded WTP and reported WTP across ROIs at the single-trial level. Each blue dot represents one participant; the black dot and error bars denote the group mean ± 1 SEM. ** p < 0.01 after FDR correction. (e) Binned scatter plots showing the positive correlation between decoded WTP and WTP reports for vmPFC (left) and SPL-IPS (right). For visualization, trials were sorted by decoded WTP and grouped into four bins (color-coded) within each participant. The black line represents the best linear fit.

We applied decoding to three ROIs that have been previously implicated in value-based decision-making (Figure 2b): vmPFC, ventral striatum and SPL-IPS (Bartra et al., 2013; Chung et al., 2025; Grabenhorst & Rolls, 2011; Harris et al., 2011; Rushworth et al., 2011). Both the vmPFC and ventral striatum have been identified as key regions for valuation across both choice tasks and continuous value estimation tasks (Bartra et al., 2013; Chung et al., 2025; Grabenhorst & Rolls, 2011; Harris et al., 2011; Kahnt, 2018; Padoa-Schioppa & Assad, 2006; Rich & Wallis, 2016; Rushworth et al., 2011). A recent meta-analysis focusing on WTP, as used in our study, confirmed that WTP is positively correlated with fMRI BOLD activity in both regions (Newton-Fenner et al., 2023). Although the SPL-IPS is not typically implicated in continuous value estimation tasks, it has long been considered relevant for representing action value (Hare et al., 2011; Louie et al., 2011; Platt & Glimcher, 1999; Roitman & Shadlen, 2002) and processing external uncertainty such as outcome probability (Chung et al., 2025; Grubb et al., 2016; Huettel et al., 2005) , in choice tasks. We found that subjective value decoded from the voxel activity (decoded WTP) significantly correlated with reported WTP in vmPFC (r = 0.120 ± 0.026, p = 0.001; binned r = 0.572, p < 0.001) and SPL-IPS (r = 0.185 ± 0.035, p = 0.001; binned r = 0.741, p < 0.001), revealing that subjective value can be decoded at a single-trial level by our decoder in these regions (Figure 2c-e; Supplementary Figure 4). Decoding performance was at a chance level in ventral striatum (Figure 2d). Because uncertainty estimates are only interpretable when the underlying value signal can be reliably decoded, we restricted subsequent analyses on decoded uncertainty to vmPFC and SPL–IPS.

### Decoded uncertainty predicts behavioral uncertainty reports

Crucially, if the decoded distributions reflect the probabilistic value representations that guide behaviors, the uncertainty of the decoded distributions should predict uncertainty reports (range), and valuation precision reflected in WTP error.

Here, we aimed to examine whether trial-by-trial variability in neural representations within the same item predicts behavioral uncertainty. To this end, we fit an LMM to predict behaviorally reported range using decoded uncertainty as the predictor while accounting for the distance to boundary, item identity, individual differences, and their interaction. We found a significant positive effect of decoded uncertainty on behaviorally reported range in vmPFC (β = 0.029 ± 0.008, p = 0.001; Figure 3a), indicating that greater neural uncertainty of value in this area was associated with higher behaviorally reported uncertainty. Individual- and group-level scatter plots further illustrated this positive correlation at the trial level in vmPFC (Figure 3b). We did not find such a relationship between decoded uncertainty and reported range in SPL-IPS (β = 0.006 ± 0.008, p = 0.470; Figure 3a).

**Figure 3.**
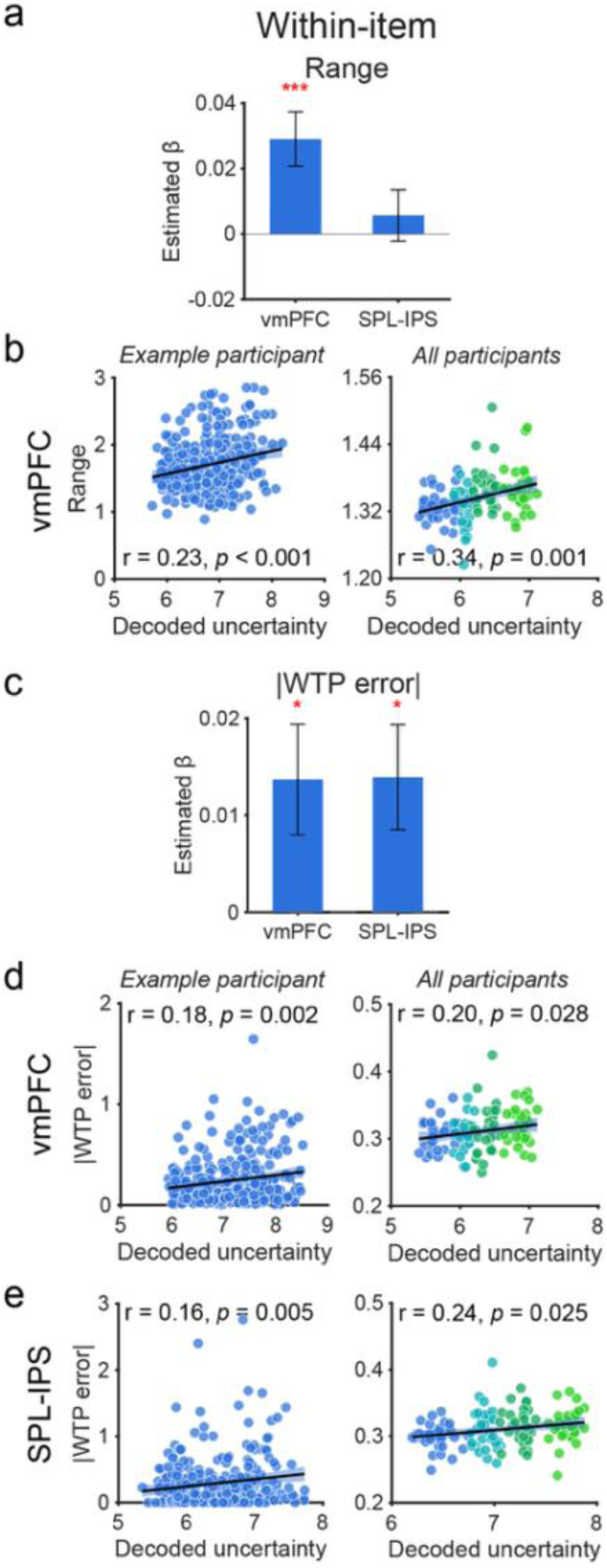
Decoded value uncertainty predicts uncertainty reports and the magnitude of WTP error. (a) Linear mixed-effects model (LMM) estimated β for the effect of decoded uncertainty on within-item trial level range report, after controlling the effect of boundary distance, the identity of the participant, item and an interaction term between item and participant. Error bars denote ±1 SEM. *** p < 0.001 after FDR correction. (b) Visualization of the positive relationship between decoded uncertainty and reported range in vmPFC for an example participant (left) and at the group level (right). Left: each dot represents a single trial from one participant. Right: each dot represents one bin of trials from a participant (four bins per participant, color-coded), sorted by decoded uncertainty. The black line represents the best-fitting linear regression. (c) LMM estimated β for the effect of decoded uncertainty on trial level WTP error across ROIs, with the same covariates as in (a). *p < 0.05, **p < 0.01 after FDR correction. (d-e) Visualization of the positive relationship between decoded uncertainty and WTP error in vmPFC (d) and SPL&IPS (e), following the same format as in (b).

We next asked whether decoded uncertainty predicted valuation variability, as indexed by trial-by-trial absolute WTP error. By LMM models, we found that decoded uncertainty significantly predicted the magnitude of WTP error in both vmPFC (β = 0.014 ± 0.006, p = 0.016; Figure 3c) and SPL-IPS (β = 0.014 ± 0.005, p = 0.016) while accounting for the distance to boundary, item identity, individual differences, and their interaction. Individual- and group-level scatter plots further illustrated this relationship at the trial level in both vmPFC (Figure 3d) and SPL-IPS (Figure 3e). Together, these results demonstrate that decoded neural uncertainty predicts within-item, trial-by-trial, behavioral uncertainty in valuation, as reflected in both explicit uncertainty reports (vmPFC) and variability in WTP (vmPFC and SPL-IPS).

While within-item fluctuations in valuation and uncertainty provide strong evidence for probabilistic coding that tracks moment-to-moment uncertainty in valuation, neural representations may also reflect more stable, item-specific uncertainty. We therefore examined whether decoded uncertainty captures not only within-item trial-level fluctuations but also stable differences across items. For each participant, we averaged decoded uncertainty or reported range across repeated presentations of the same item. For both vmPFC and SPL-IPS, we found that items with higher decoded uncertainty were reported with a larger range while controlling for the boundary distance and individual differences (vmPFC: β = 0.104 ± 0.036, p = 0.004; SPL-IPS: β = 0.105 ± 0.034, p = 0.004; Figure 4a&b). We conducted a similar analysis to investigate the relationship between item-wise decoded uncertainty and item-wise WTP variability. Decoded uncertainty predicted item-wise WTP variability only in SPL-IPS when boundary distance was not included in the regression (Supplementary Figure 5), and the effect was not significant when boundary distance was included in the LMM (p > 0.583). Together, these results indicate that decoded uncertainty captures stable, item-specific valuation uncertainty as reflected in explicit uncertainty reports in both vmPFC and parietal cortex. Item-wise WTP variability, may be a composite measure reflecting not only the uncertainty in the underlying value representation but also additional variability arising from the value reporting process.

**Figure 4.**
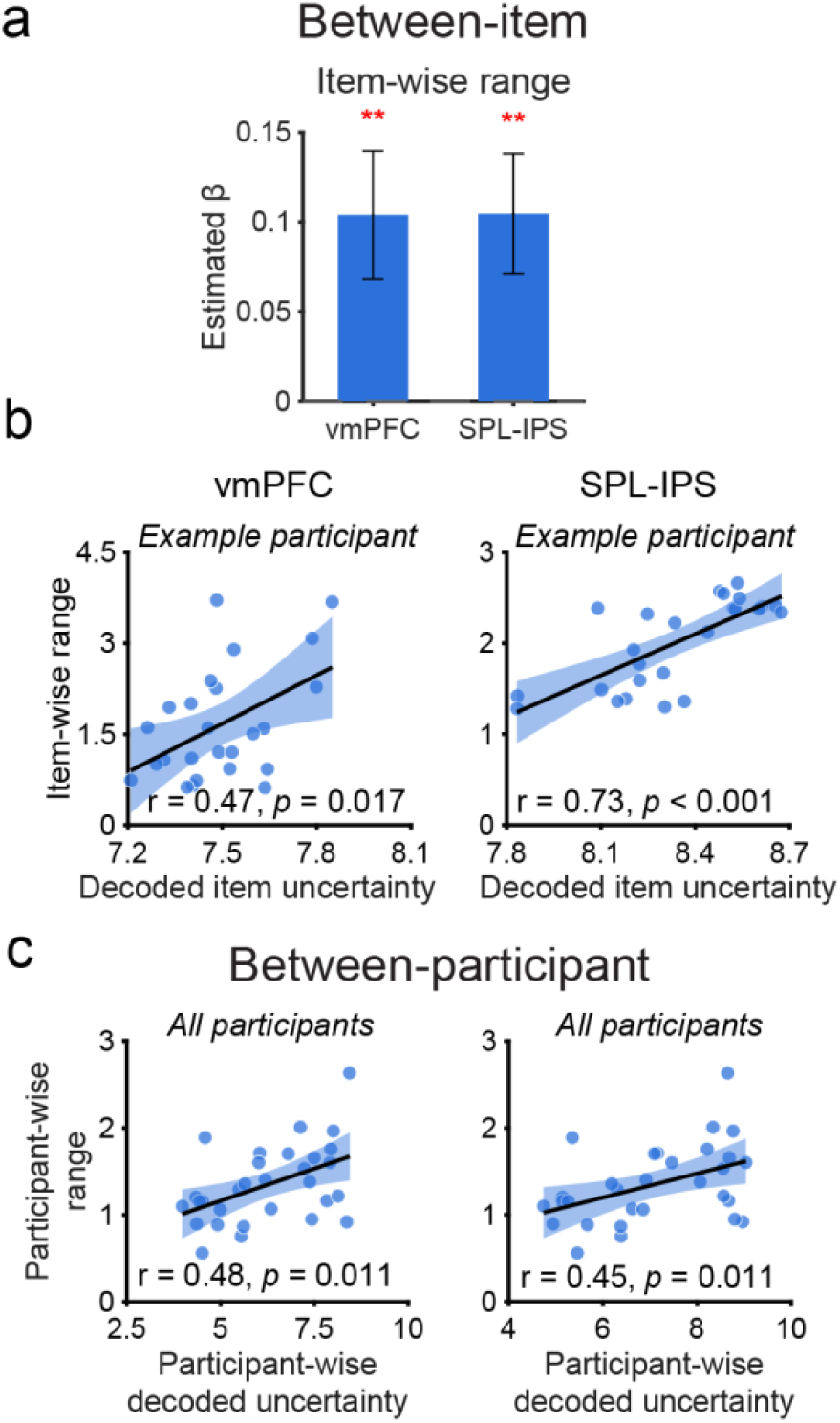
Decoded value uncertainty predicts item- and individual-level uncertainty reports. (a) LMM estimated β for the effect of decoded uncertainty on item-averaged range report across ROIs, controlling for item WTP distance to scale boundary and random effects of participant. Error bars denote ±1 SEM. **p < 0.01 after FDR correction. (b) Correlation between item-averaged decoded uncertainty and item-averaged range report in vmPFC (left) and SPL-IPS (right) for an example participant. Each dot represents one food item. The black line represents the best linear fit. (c) Correlation between participant-averaged decoded uncertainty and participant-averaged range report in vmPFC (left) and SPL-IPS (right). Each dot represents one participant. The black line represents the best linear fit.

### Individual differences in neural uncertainty predict behavioral valuation uncertainty

Finally, we examined whether decoded uncertainty from neural activity can explain individual differences in behavioral uncertainty reports. By averaging the decoded uncertainty, or reported range, across all items for each participant, we found that participants reporting higher uncertainty (larger range) behaviorally were associated with higher overall decoded uncertainty in both vmPFC (r = 0.48, p = 0.011; Figure 4c) and SPL-IPS (r = 0.45, p = 0.011). These results demonstrate a linkage between the precision of value neural representation in these brain regions and the individual differences in subjective uncertainty of value estimation.

## Discussion

In this study, we investigated whether subjective value is represented probabilistically, allowing individuals to have metacognitive access to uncertainty in their value estimates. We found converging behavioral and neural evidence in support of this hypothesis. We first show behaviorally that uncertainty reports tracked trial-level variability in valuation, quantified as

WTP error, demonstrating that humans have metacognitive access to trial-by-trial fluctuations in the quality of valuation. Building on this, we decoded probability distributions over value from neural population activity in vmPFC and SPL-IPS and showed that decoded neural uncertainty predicts behavioral uncertainty. Together, these findings provide evidence that subjective value is represented as a probability distribution in the human brain, offering a neural basis for metacognitive access to uncertainty in value estimation.

Our behavioral findings provide evidence for metacognitive sensitivity in subjective value estimation. Previous studies have examined confidence in value-based decisions, often emphasizing the characteristic U-shaped relationship between confidence and the value rating scale, which we also observed as an inverted-U pattern for uncertainty reports. This pattern arises from the bounded nature of the rating scale, which constrains the underlying belief distribution and can serve as a proxy for confidence or certainty. Importantly, prior work has leveraged this signature to investigate the neural basis of confidence in valuation. Building on these findings, our analysis goes beyond this boundary-driven effect. After accounting for scale constraints, we focused on trial-by-trial variability in valuation within the same item. Such variability likely reflects internal neural noise in value representations, and we show that it is systematically tracked by participants’ uncertainty reports. This result implies that, on each trial, individuals maintain an internal estimate of uncertainty alongside value, consistent with an underlying probabilistic representation of value.

Our approach provides a window into metacognitive processes in valuation. In perceptual and memory domains, metacognition in estimation is often assessed by relating confidence or uncertainty judgments to objective estimation error or variability of estimation (Geurts et al., 2022; Honig et al., 2020; Li et al., 2021; Van den Berg et al., 2017). Here we show that a similar relationship can be established in subjective value estimation: uncertainty reports track fluctuations in valuation estimates even in the absence of an explicit ground truth. These findings suggest that metacognitive representations of uncertainty may operate under similar principles across perceptual, memory, and value-based domains.

The ability to monitor trial-by-trial uncertainty in valuation implies a neural code that represents more than a point estimate of value. A candidate for such a representation is the framework of probabilistic population codes. Originally developed to explain perception, probabilistic population coding theory posits that neural populations encode both stimulus estimates and their associated uncertainty by representing full probability distributions over stimulus space (Beck et al., 2008; Földiák, 1993; Jazayeri & Movshon, 2006; Ma et al., 2006; Sanger, 1996; Zemel et al., 1998). Under this framework, the brain leverages the knowledge of the generative process governing neural responses, allowing variability (neural noise) in population activity to be interpreted, via Bayesian inference, as a probability distribution over the underlying stimulus.

Empirical support for probabilistic population code has been observed in neurophysiological recordings from non-human primates, where population activity in visual cortex reflects both sensory estimates and their uncertainty (Berens et al., 2012; Graf et al., 2011; Walker et al., 2020). More recently, computational neuroimaging approaches have extended this framework to human fMRI, showing that probabilistic information can be decoded from population activity in the human brain to predict uncertainty and confidence reports in visual perception (Geurts et al., 2022; Van Bergen et al., 2015) and working memory (Li et al., 2021, 2025). Building on this line of work, the present findings suggest that similar computational principles extend to the domain of value, with neural populations encoding probabilistic representations that support both valuation and metacognitive access to uncertainty.

Our results also highlight a shift in how value representations are characterized in the human brain. Much of the existing literature has focused on decoding a single estimate of value using a simple linear decoder, and extracting metrics such as representational similarity or decoding accuracy by aggregating multiple trials (Kahnt, 2018; LoFaro et al., 2025; Suzuki et al., 2017). In contrast, our approach leverages trial-by-trial multivariate variability to recover full probability distributions over value, revealing that behaviorally meaningful neural noise as an essential part of the neural code for subjective value, and can be utilized at the single-trial level for value decoding. Notably, this probabilistic perspective resonates with recent findings in animal studies of reward learning, where value representations have been found to follow a distributional code that captures the full range of possible outcomes rather than a single expected value in ventral tegmental area (Dabney et al., 2020; Lowet et al., 2020, 2025; Muller et al., 2024) and anterior cingulate cortex (Muller et al., 2024).

A recent study similarly found evidence for a probabilistic neural code for value in vmPFC (Bouc et al., 2026). Together, Bouc et al. and our study provide converging evidence for probabilistic representations of value in the human brain. At the same time, there exist key differences between the two studies. We repeated each item 4 (Experiment 1) and 12 times (Experiment 2), allowing us to explicitly examine trial-by-trial fluctuations (WTP error) in valuation within the same item. In addition, our analyses included item and participants’ identity as covariates. These approaches uniquely enabled us to directly relate both behavioral uncertainty report and neural uncertainty to within-item variability in valuation, isolating fluctuations that reflect internal noise rather than item-specific properties. This highlights the role of metacognition in tracking moment-to-moment changes in valuation.

Our results that vmPFC involved in value and uncertainty processing are consistent with the findings that vmPFC and adjacent lateral orbitofrontal cortex automatically encode confidence and uncertainty signals alongside subjective value even when no explicit report is required (Clairis & Pessiglione, 2022; Lebreton et al., 2015; Lopez-Persem et al., 2020). The previous studies primarily identified such signals using univariate analyses by exploiting the characteristic U-shaped relationship between confidence and value induced by scale boundaries. In contrast, our multivariate decoding approach recovers trial-by-trial distributions over value and extracts their associated uncertainty, providing a complementary perspective on how uncertainty is represented. Uncertainty or confidence in vmPFC may be encoded in multiple formats. Populations that encode value would contain uncertainty information through probabilistic representations that can be revealed through decoding, while some neural populations may represent overall confidence level as a scalar signal detectable with univariate analyses. Importantly, we found that decoded uncertainty from vmPFC predicted explicit uncertainty reports at the trial, item, and individual levels, extending this prior work by demonstrating that the value uncertainty in vmPFC is not only present implicitly but is also accessible to conscious metacognitive report across multiple scales.

Beyond vmPFC, decoded uncertainty from SPL-IPS predicted trial-level WTP error as well as item- and individual-level uncertainty reports, suggesting that parietal cortex encodes valuation uncertainty in a format that captures both moment-to-moment fluctuations in behavioral precision and stable item- and individual-specific properties. Unlike vmPFC, however, SPL-IPS uncertainty did not predict trial-level explicit uncertainty reports, suggesting that the parietal signal influences behavioral precision without being directly accessible to conscious introspection. This dissociation is consistent with the established role of parietal cortex in value-based action selection (Platt & Glimcher, 1999; Rushworth et al., 2011) and in encoding the acuity of numerical magnitude representations (Barretto-García et al., 2023; de Hollander et al., 2025), processes that may shape valuation precision implicitly rather than through metacognitive awareness.

In this study, we broadly decomposed uncertainty into three components (Brus et al., 2021; Fleming, 2024; Pouget et al., 2016). At the within-item (trial) level, transient fluctuations in attentional engagement, working memory resources, and intrinsic neural noise during valuation may introduce moment-to-moment variability in representational precision. At the item level, stable uncertainty likely reflects the intrinsic properties of each option, such as attribute ambiguity, and the individual’s limited prior experience with it, both of which complicate precise value computation. At the individual level, reliable between-subject differences in both decoded neural precision and explicit uncertainty reports suggest that the fidelity of value representations is also a stable trait, potentially linked to individual differences in the efficiency of value construction and its underlying neural circuitry. Notably, we find that decoded uncertainty relates to all three levels, suggesting that this neural representation is, to some extent, source-invariant. A similar pattern was observed in our previous work on visual working memory (Li et al., 2021), where we decomposed behavioral uncertainty at different levels, and found that neurally decoded uncertainty predicted not only trial-by-trial fluctuations in memory uncertainty reports but also stable individual differences in memory precision. Together, these findings point to a probabilistic neural code as a common computational mechanism that operates across multiple scales and domains to represent uncertainty.

## Methods

### Participants

A total of 44 human participants (17 males; 19–32 years old) participated in the study. All completed the behavioral experiment (Experiment 1), which included a subjective value estimation task and a 2AFC choice task. A subset of 30 participants (13 males; 20–31 years old) subsequently underwent fMRI scanning while only performing the subjective value estimation task. All participants reported normal or corrected-to-normal vision, and reported no history of known neurological, eating-related, or psychiatric disorders. They were naive to the purpose of the study and provided written informed consent prior to participation. All procedures were approved by the Institutional Review Board (IRB) at The Ohio State University.

### Apparatus and Stimuli

Visual stimuli were generated using MATLAB (MathWorks, Natick, MA) with the Psychophysics Toolbox (Brainard & Vision, 1997), with head position stabilized using a chin rest. A white fixation dot (diameter 0.25°) was present at the center of the screen throughout the experiment. Stimuli consisted of 30 photographs of commercially available food items presented on a gray background, each subtending 7° × 7° of visual angle. Value and uncertainty reports were made on a continuous $0–$5 scale subtending 9° of visual angle, using a four-key interface allowing cursor movement at coarse and fine speeds, confirmed with a fifth key.

### Experiment 1 - Behavioral Experiments

**Subjective value estimation task.** Participants fasted for 3 h before the experiment and completed a subjective value estimation task to elicit both subjective value and uncertainty estimates for each food item (common salty and sweet snack foods), using an incentive-compatible procedure known as the Becker–DeGroot–Marschak (BDM) mechanism (Becker et al., 1964). Each trial began with the onset of a food image presented for 6 s. Participants then had a 10 s response window in which they made two sequential reports on a horizontal rating scale ranging from $0 to $5, with scale orientation alternating every two blocks. They first indicated their willingness to pay (WTP) for the item, then reported an uncertainty range defined by the distance between the minimum price below which they would certainly purchase and the maximum price above which they would certainly not purchase the item. Participants were informed that both the WTP and range reports were subjective, and they were allowed to set the minimum or maximum price exactly at their reported WTP. The cursor was initialized at a random position on the scale for the WTP report, and the order of minimum and maximum range reports was randomized across trials, with each cursor initialized at a random position between the WTP and the corresponding scale boundary ($0 or $5). Following the response window, a 1 s feedback display indicated whether the trial was completed in time. The intertrial interval was pseudo randomly drawn from 1, 1.5, 2, 2.5, or 3 s. Each participant completed four blocks of 30 trials, with each of the 30 food items appearing once per block (120 trials total).

At the end of the experiment, one trial was randomly selected and the participant had the opportunity to buy this item by means of an auction administered according to the BDM procedure (Becker et al., 1964). Specifically, participants received an endowment of $5 and the computer drew a market price for the selected auction food item from a uniform distribution ($0–$5). If the participant’s WTP exceeded the market price, the participant purchased the item at the market price using the endowed $5; if the WTP was below the market price, no transaction occurred. Participants were required to remain in the lab for an additional 30 minutes following the experiment, during which the only food they were permitted to consume was the item purchased in the auction, if any. This BDM procedure ensured that participants had a financial incentive to report their true subjective value (willingness-to-pay).

**2AFC Task.** Participants next completed a two-alternative forced-choice (2AFC) task in which food stimuli were selected individually based on item-level WTP estimates (mean WTP across item repetition) from the subjective value estimation task. Five items with WTP estimates nearest to the scale boundaries were excluded, yielding 25 retained items, which were used in the subsequent fMRI experiment. For each retained item, four comparison items were drawn from the remaining 29 snacks with WTP differences of approximately $0.25, $0.50, $0.75, and $1.00, respectively. Each trial began with the simultaneous onset of two food images presented in the upper and lower visual fields for 4 s, during which participants indicated their preferred item via keypress. The selected image was then highlighted with a red frame, followed by a 1 s feedback display indicating whether the response was made in time. Each pair was presented four times with image position counterbalanced across trials, yielding 400 trials completed across five blocks.

### Experiment 2 - fMRI Experiment

The fMRI experiment followed the same procedures as the subjective value estimation task in the behavioral experiment, with four exceptions. First, the stimulus set was restricted to 25 retained items, selected by excluding the five snacks with WTP estimates nearest to the scale boundaries. Second, scale orientation was counterbalanced across the two scanning sessions rather than alternating every two blocks, with the session-to-orientation assignment randomized across participants. Third, the feedback display duration was extended to 2 s. Fourth, the intertrial interval was pseudo randomly drawn from 4, 6, 8, 10, or 12 s to allow adequate sampling of the hemodynamic response. Each participant completed 25 trials per block across 10–12 blocks in two scanning sessions (250–300 trials total). A BDM auction was held at the end of each scanning session, following the same procedure as described above.

### Behavioral Data Analysis

To quantify subjective value at the item level, we averaged each participant’s WTP reports across trials for each snack item:

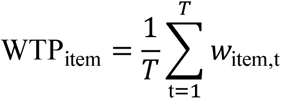

where 𝑤_item,t_ denotes the WTP report for a given item on trial 𝑡, and 𝑇 is the total number of trials for that item. Trial-level WTP error, reflecting the consistency of individual valuation reports relative to each participant’s stable preference for a given snack, was defined as the absolute difference between a single-trial WTP report and the corresponding item-wise mean:

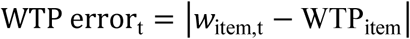

Because WTP reports near the scale boundaries ($0 or $5) are inherently constrained, we computed each trial’s distance to the nearest boundary as a control variable DTB_𝑡_= min(𝑤_item,𝑡_, 5 − 𝑤_item,𝑡_).We first examined the zero-order correlation between Range and WTP error, then fit a linear mixed-effects model (LMM) to test whether range independently predicted valuation consistency after controlling for boundary distance, with crossed random effects for participants (𝑢_𝑖_), items (𝑣_𝑗_) and their interaction (𝑤_𝑖𝑗_ ), where subscripts 𝑖, 𝑗 and 𝑡 index participants, items, and trials, respectively:

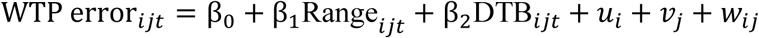

A significant positive 𝛽_1_ would indicate that wider reported uncertainty ranges predict greater trial-to-trial variability in WTP, independently of boundary proximity.

To quantify uncertainty associated with each choice pair in the 2AFC Task, we derived two complementary measures from subjective value estimation task. Item-wise uncertainty was defined as the mean reported range across trials:

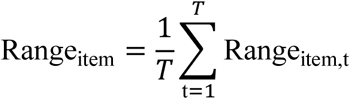

Item-wise WTP variability was defined as the standard deviation of WTP reports across trials (repetitions) for an item.

For each choice trial, pair-level measures were then derived from these item-wise estimates: Total Range (TR) and Total Variability (TV) were computed as the sum of the two items’ and Range_item_ and VarWTP_item_ measures, respectively; value difference (VD) was defined as the absolute WTP difference between the two items, and value sum (VS) as the sum of their WTP estimates.

To examine whether these uncertainty measures predicted choice behavior, we fit generalized linear mixed-effects models (GLMMs) with a binomial distribution and logit link, each including a random intercept for participants 𝑢_𝑖_. Two models tested whether uncertainty predicted the accuracy of value-based choices, with choice accuracy (1 if the higher-valued item was chosen, 0 otherwise) as the outcome:

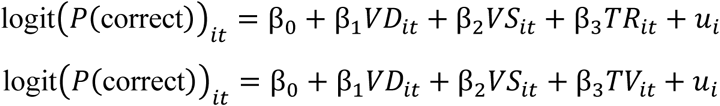

where subscripts 𝑖 and 𝑡 index participants and trials, respectively, and β_3_ reflects the predictive contribution of uncertainty.

Furthermore, to examine whether valuation uncertainty influenced decision deliberation, we tested whether TR and TV each predicted response times (RT) in the choice task, using separate LMMs with VD and VS as covariates and a random intercept for participants:

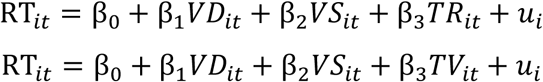

A significant positive β_3_ would indicate that higher valuation uncertainty is associated with longer deliberation times, independent of value difference and overall value.

The same analyses were applied to behavioral data collected during the fMRI experiment, using identical variable definitions. Notably, no choice task was performed following the fMRI experiment, so choice-task-related analysis were not included.

### MRI Data Acquisition

MRI data were collected using a 3T Siemens MAGNETOM XR scanner with a 32-channel phased-array head receiver coil at the Center for Cognitive and Behavioral Brain Imaging (CCBBI) at The Ohio State University. Blood-oxygen-level-dependent (BOLD) signals were measured with a multiband gradient-echo echo-planar imaging sequence (TE: 28 ms; TR: 2,000 ms; FOV: 220 × 220 mm²; matrix: 100 × 100; flip angle: 76°; slice thickness: 2.2 mm; gap: 0 mm; number of slices: 69; multiband acceleration factor: 3; phase partial Fourier: 7/8; slice orientation: axial, interleaved). Each participant completed two functional sessions, with a total of 10–12 runs across sessions (356 volumes per run; run duration: ∼12 min). Spin-echo EPI field maps were additionally acquired in both anterior-to-posterior and posterior-to-anterior phase-encoding directions (TE: 61 ms; TR: 7,120 ms; 3 volumes each) to correct for susceptibility-induced geometric distortions.

A high-resolution T1-weighted 3D structural dataset (3D MPRAGE; 0.8 × 0.8 × 0.8 mm³ resolution; TR: 2,400 ms; TE: 2.24 ms; TI: 1,060 ms; FOV: 256 × 240 mm²; flip angle: 8°; number of slices: 208; GRAPPA acceleration factor: 2; slice orientation: sagittal) and a T2-weighted anatomical image (3D SPACE; 0.8 × 0.8 × 0.8 mm³ resolution; TR: 3,200 ms; TE: 563 ms; number of slices: 208; GRAPPA acceleration factor: 2) were collected at the end of the first session, following the functional scans.

### MRI Data Preprocessing

Results included in this manuscript come from preprocessing performed using fMRIPrep 24.1.1. (Esteban et al., 2019; RRID:SCR_016216), which is based on Nipype 1.8.6 (Gorgolewski et al., 2011; RRID:SCR_002502).

**B0 field map preprocessing.** A total of 2 fieldmaps were found available within the input BIDS structure. A B0-nonuniformity map (or fieldmap) was estimated based on two echo-planar imaging (EPI) references with opposite phase-encoding directions using topup (Andersson et al., 2003; FSL None).

**Anatomical data preprocessing.** The T1-weighted (T1w) image was corrected for intensity non-uniformity (INU) with N4BiasFieldCorrection (Tustison et al., 2010), distributed with ANTs 2.5.3 (Avants et al., 2008; RRID:SCR_004757), and used as T1w-reference throughout the workflow. The T1w-reference was then skull-stripped with a Nipype implementation of the antsBrainExtraction.sh workflow (from ANTs), using OASIS30ANTs as target template. Brain tissue segmentation of cerebrospinal fluid (CSF), white matter (WM) and gray matter (GM) was performed on the brain-extracted T1w using fast (FSL; RRID:SCR_002823; Zhang et al., 2001). Brain surfaces were reconstructed using recon-all (FreeSurfer 7.3.2; RRID:SCR_001847; Dale et al., 1999), and the brain mask was refined with a custom variation of the method to reconcile ANTs-derived and FreeSurfer-derived segmentations of the cortical gray-matter of Mindboggle (RRID:SCR_002438; Klein et al., 2017). A T2-weighted image was used to improve pial surface refinement. Volume-based spatial normalization to standard space (MNI152NLin2009cAsym) was performed through nonlinear registration with antsRegistration (ANTs 2.5.3), using brain-extracted versions of both T1w reference and the T1w template (ICBM 152 Nonlinear Asymmetrical template version 2009c; Fonov et al., 2011; RRID:SCR_008796; TemplateFlow ID: MNI152NLin2009cAsym).

**Functional data preprocessing.** For each of the BOLD runs found per participant (across all sessions), the following preprocessing was performed. First, a reference volume and its skull-stripped version were generated using a custom methodology of fMRIPrep. Head-motion parameters with respect to the BOLD reference (transformation matrices, and six corresponding rotation and translation parameters) were estimated before any spatiotemporal filtering using mcflirt (FSL; Jenkinson et al., 2002). The estimated fieldmap was aligned with rigid-registration to the target EPI reference run and mapped onto the reference EPI. The BOLD reference was then co-registered to the T1w reference using bbregister (FreeSurfer), which implements boundary-based registration (Greve & Fischl, 2009), configured with six degrees of freedom.

Several confounding time-series were calculated based on the preprocessed BOLD: framewise displacement (FD), DVARS, and three region-wise global signals. FD and DVARS are calculated for each functional run using their implementations in Nipype (Power et al., 2014). Frames that exceeded a threshold of 0.5 mm FD or 1.5 standardized DVARS were annotated as motion outliers. Additionally, physiological regressors were extracted to allow for component-based noise correction (CompCor; Behzadi et al., 2007), including temporal (tCompCor) and anatomical (aCompCor) variants. Principal components were estimated after high-pass filtering the preprocessed BOLD time series using a discrete cosine filter with a 128 s cutoff. For each CompCor decomposition, components were retained until the cumulative explained variance exceeded 50% of the nuisance mask signal. The head-motion estimates were further expanded with temporal derivatives and quadratic terms for each (Satterthwaite et al., 2013).The BOLD time-series were resampled onto the following surfaces (FreeSurfer reconstruction nomenclature): fsaverage6, fsnative. Gridded (volumetric) resamplings were performed using nitransforms, configured with cubic B-spline interpolation. Non-gridded (surface) resamplings were performed using mri_vol2surf (FreeSurfer). Many internal operations of fMRIPrep use Nilearn 0.10.4 (Abraham et al., 2014; RRID:SCR_001362).

### Region of Interest (ROI) Definition

Regions of interest (ROIs) were defined based on prior studies implicating these areas in value-based decision-making, including the vmPFC, ventral striatum, and SPL/IPS. The vmPFC, in particular, has been consistently shown to encode the subjective value of options across species (including humans and non-human primates) and across both choice and valuation tasks (Bartra et al., 2013; Chung et al., 2025; Grabenhorst & Rolls, 2011; Harris et al., 2011; Kahnt, 2018; Padoa-Schioppa & Assad, 2006;, 2016; Rushworth et al., 2011). The vmPFC mask was defined by combining regions identified by Rolls et al. (Rolls et al., 2022), including Area 10 ventromedial prefrontal regions (10r, 10v, 10d, 9m), subgenual cingulate cortex (area 25), and pregenual anterior cingulate cortex (s32, a24, p24, p32, d32), with lateral and medial orbitofrontal cortex regions: lateral OFC (47s, 47l, a47r, p47r, 47m) and medial OFC (11l, 13l, OFC, pOFC). Similarly, the ventral striatum has been shown to encode value and plays a key role in value-based decision-making (Bartra et al., 2013; Chung et al., 2017, 2025; Harris et al., 2011; Kahnt, 2018; Knutson et al., 2001). The ventral striatum ROI was defined using the HCP-MMP extended atlas (Glasser et al., 2016; Huang et al., 2022), comprising the bilateral putamen. The SPL/IPS has been implicated in representing action value (Hare et al., 2011; Louie et al., 2011; Roitman & Shadlen, 2002) and in processing external uncertainty, such as outcome probability, during choice tasks. The SPL/IPS is also involved in numerosity representation (Harvey et al., 2013; Harvey & Dumoulin, 2017; Nieder & Dehaene, 2009) and recent work suggests that these representations in the SPL/IPS may be recruited for value-based decision-making (Barretto-García et al., 2023; de Hollander et al., 2025). SPL-IPS mask comprising 10 areas from the HCP-MMP atlas (Glasser et al., 2016): LIPv, LIPd, VIP, AIP, MIP, 7PC, 7AL, 7Am, 7PL, and 7Pm, spanning the medial bank of the intraparietal sulcus and adjacent superior parietal cortex. All ROIs were defined in MNI152 standard space and cortical regions were additionally visualized on surface maps (Figure 2b).

### Single Trial Beta Estimates

We used the GLMSingle matlab package (Prince et al., 2022) to obtain single-trial BOLD estimates. Briefly, GLMsingle optimizes three components via cross-validation:: a library of hemodynamic response functions to identify the best-fitting HRF for each voxel; L2-regularization parameters to shrink single-trial estimates and address the issue of correlated single-trial regressors (Mumford et al., 2012); and GLMDenoise regressors (Kay et al., 2013), which are nuisance regressors derived from principal components of voxels unrelated to the experimental paradigm, with the optimal number determined via cross-validation

As input to GLMSingle, we modeled each snack food presentation as a separate trial regressor, with a duration of 6 s convolved with the hemodynamic response function, to obtain trial-wise beta estimates. The response window and the feedback display were each modeled as separate regressors of no interest, collapsed across all trials. No additional confound regressors (e.g., motion parameters, RETROICOR, or aCompCor regressors) were included beyond the GLMDenoise confound regressors, as the fMRIPrep preprocessed MNI152-space BOLD data were used directly as input.

### fMRI Decoding

**Generative model.** We decoded subjective value from BOLD responses using a generative model (TAFKAP; Van Bergen et al., 2015; Van Bergen & Jehee, 2021). This Bayesian decoding approach allows us to concurrently read out estimated value and decoding uncertainty from the same decoded probability distribution. In this generative model, the multivariate voxel response given the subjective value ($0–$5) was modeled as a multivariate normal distribution. The mean response of each voxel given a stimulus was determined by its tuning function (voxel response as a function of subjective value). The voxel tuning function was approximated by a weighted sum of *K* = 8 Gaussian basis functions that evenly tiled the value space:

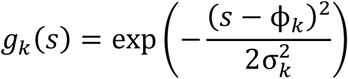

where ϕ_𝑘_ is the center of the *k*-th basis function, and the *K* centers ϕ_𝑘_ are spaced evenly across the $0–$5 value space. The response of the 𝑖-th voxel given a stimulus 𝑠 is then modeled as:

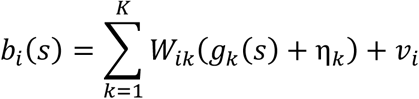

where 𝑊 is a weighting matrix that determines the weights of each basis function for each voxel. Because each voxel reflects the aggregate activity of many neurons with potentially diverse tuning properties, its response profile with respect to value may not follow a simple parametric form. The implementation here allows each voxel to exhibit flexible tuning functions with respect to value, without imposing strong parametric constraints. In the model, two sources of variability are considered. First, η_𝑘_ is the noise specific to each basis function. This noise was carried over into each voxel by the weighting matrix 𝑊, modeling noise shared across voxels with similar tuning functions. The model assumed that η follows a zero-mean normal distribution whose covariance matrix is a constant noise magnitude multiplied with an identity matrix η ∼ 𝒩(0, σ^2^𝐼). Second, 𝑣_𝑖_ represents the noise specific to each voxel. The model assumed that the voxel-wise noise follows a zero-mean normal distribution 𝑣 ∼ 𝒩(0, Σ). The covariance matrix of this distribution is approximated by a rank-one covariance matrix plus a diagonal matrix:

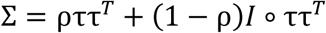

where ∘ represents the Hadamard product, i.e., the element-wise product between two matrices. Thus, based on this generative process, the theoretical covariance matrix of the multivariate response of the voxels given a stimulus 𝑠 is:

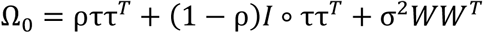

The first two terms of the theoretical covariance matrix consider a simple form of covariance as a weighted sum between a diagonal matrix (where τ is a vector representing the standard deviation of the noise of each voxel) and a rank-one covariance matrix. The last term captures the covariance depending on the tuning functions of the voxels (i.e., voxels selective for similar values have higher covariance; see derivations in Van Bergen et al., 2015). In addition to the theoretical covariance matrix, the model also considered the empirical sample covariance:

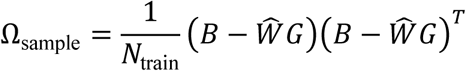

where 𝐵 is the training data and 𝐺 is the response of the basis functions given the training set stimuli. Thus, for each training dataset, we assumed that the voxel activity pattern followed a multivariate normal distribution:

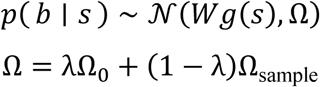

When the number of variables (voxels) is larger than the number of observations (trials), the sample covariance is not invertible. To ensure an invertible and stable estimation of the covariance matrix, the covariance matrix was modeled as the sample covariance matrix shrunk to a target covariance matrix, the theoretical covariance matrix Ω_0_. The degree of shrinkage is determined by a free parameter λ (see details in Van Bergen & Jehee, 2021).

**Training and testing.** For each voxel, we used the single-trial beta estimates obtained from GLMSingle, z-normalized within each run, as input to the model in the main text. All voxels within each ROI were included in the main analysis. Model parameters were estimated for each participant in a leave one run out cross-validation procedure. That is, the model’s free parameters were first estimated from the data of all but one run, and the remaining run was used to decode posterior distributions on a trial-by-trial basis. Each run was used as a test set once. For each trial in the test set, the posterior distribution over subjective value was subsequently computed, conditioned on the fitted model parameters, using Bayes’ rule:

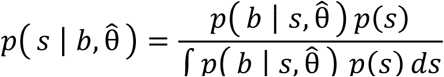

where the prior 𝑝(𝑠) was flat over the $0–$5 value space, reflecting the uniform distribution of value levels in the experiment, and the normalizing constant was estimated numerically. We took the mean of the decoded posterior distribution as the estimated subjective value and its entropy as a measure of uncertainty.

### Statistical Analysis

To examine whether decoded uncertainty predicted trial-level range report and WTP deviation, independently of scale boundary effects, we fit separate LMMs for each ROI with range report and WTP error as the respective outcome variables, decoded uncertainty and boundary distance as fixed effects, with crossed random effects for participants (𝑢_𝑖_), items (𝑣_𝑗_) and their interaction (𝑤_𝑖𝑗_ ), where subscripts 𝑖, 𝑗 𝑎𝑛𝑑 𝑡 index participants, items, and trials, respectively:

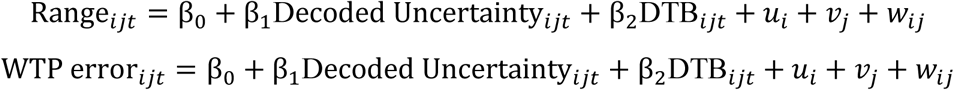

For the Range model, a significant positive 𝛽_1_ would indicate that higher decoded uncertainty is associated with wider reported uncertainty ranges. For the WTP error model, a significant positive 𝛽_1_indicates that higher decoded uncertainty predicts greater trial-to-trial variability in WTP, independently of boundary proximity. Benjamini–Hochberg false discovery rate (FDR) correction was applied across ROIs for all analyses.

For snack-level analyses, decoded uncertainty was averaged across trials within each snack item as Decoded Uncertainty_𝑖,𝑖𝑡𝑒𝑚_. Boundary distance was computed from each item’s true WTP value, as DTB_item_ = min(𝑤_item_, 5 − 𝑤_item_). We then examined whether decoded uncertainty predicted snack-level range report and WTP variability, with a random intercept for participants (𝑢_𝑖_), where 𝑖 and 𝑖𝑡𝑒𝑚 index participants and items, respectively:

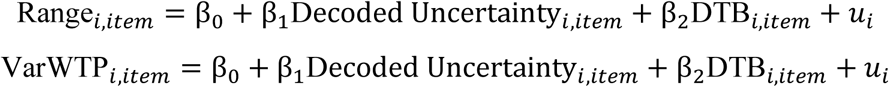

A significant positive β_1_ would indicate that higher decoded snack uncertainty is associated with wider snack uncertainty ranges and greater WTP variability across snack items, independently of boundary proximity. FDR correction was applied across ROIs for all analyses.

All correlations between decoded variables and corresponding behavioral measures were computed using Pearson correlations. At the group-level (Figure 2d), individual correlation coefficients were Fisher z-transformed before averaging across participants. Group-level significance was assessed using a bootstrap test (2,000 iterations, two-tailed) on the mean z-transformed correlation. Group-level mean correlations were back-transformed to correlation coefficient (r) for visualization and reporting. In addition to individual participant correlations, we present binned scatter plots for visualization purposes (Figures 2e&3). For each participant, trials were sorted by the x-axis variable and divided into four equal-sized bins. For each bin, we computed the mean value of the variable on the y-axis across trials within that bin. Before pooling the data across participants, both x- and y-axis values were mean-centered within each participant to remove individual baseline differences, enabling visualization of within-subject covariation. The grand mean was then added back to preserve interpretability of the absolute scale. Significance was assessed using a permutation test (2,000 permutations, two-tailed) on the pooled binned data. For figures displaying decoded WTP (Figure 2e), both x- and y-axis values were subsequently normalized to the [0, 5] range to match the scale of the original WTP reports and improve interpretability. For the binned scatter plots related to decoded uncertainty (Figure 3), we regressed out the effects of boundary distance, as well as participant and item random effects, from both x- and y-axis variables. FDR correction was applied across ROIs for all analyses.

## Acknowledgements

Author contributions

H.-H.L. and H.-K.C. conceived the study. All authors designed the experiments. H.L. collected the data and performed the analyses with input from H.-H.L. All authors contributed to writing and editing the manuscript. H.-H.L. supervised the project. All authors approved the final manuscript.

## Supplemental information

**Supplementary Figure 1.**
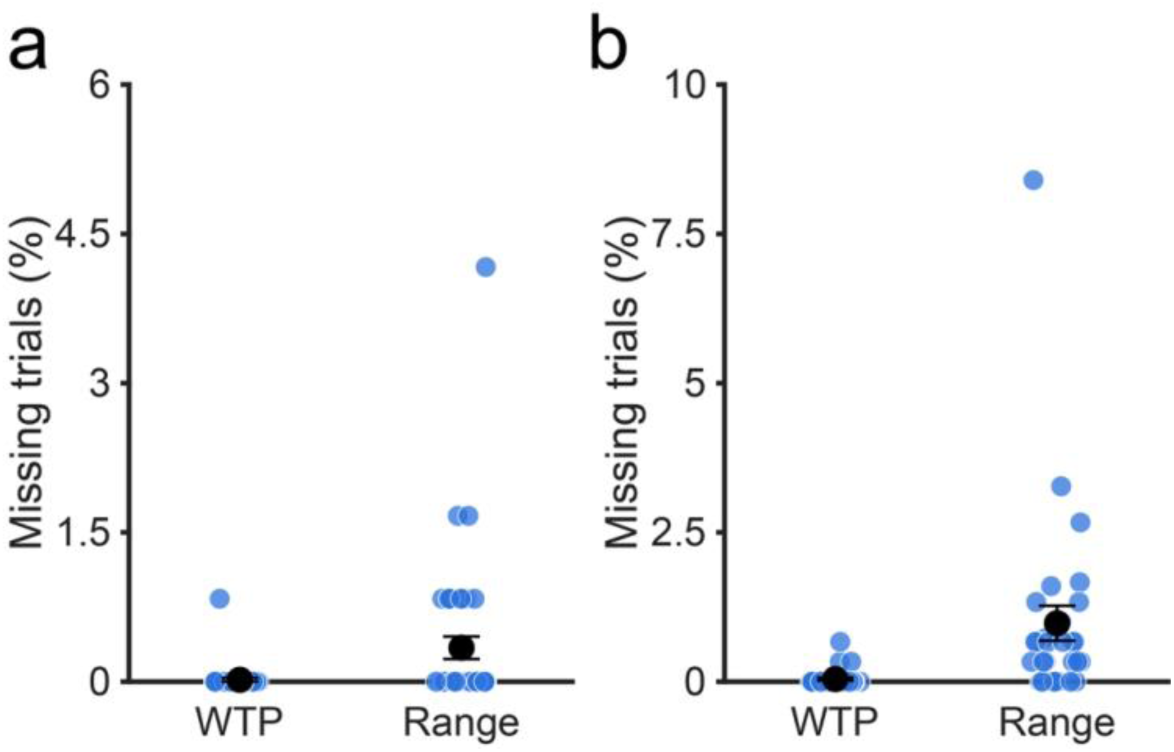
Behavioral report quality across experiments. Missing trial rates for WTP and range reports across participants in the behavioral (a; WTP: 0.02 ± 0.02%; Range: 0.34 ± 0.11%) and fMRI (b; WTP: 0.04 ± 0.03%; Range: 0.98 ± 0.29%) experiments, indicating high task compliance. The black dot and error bars denote the group mean ± 1 SEM.

**Supplementary Figure 2.**
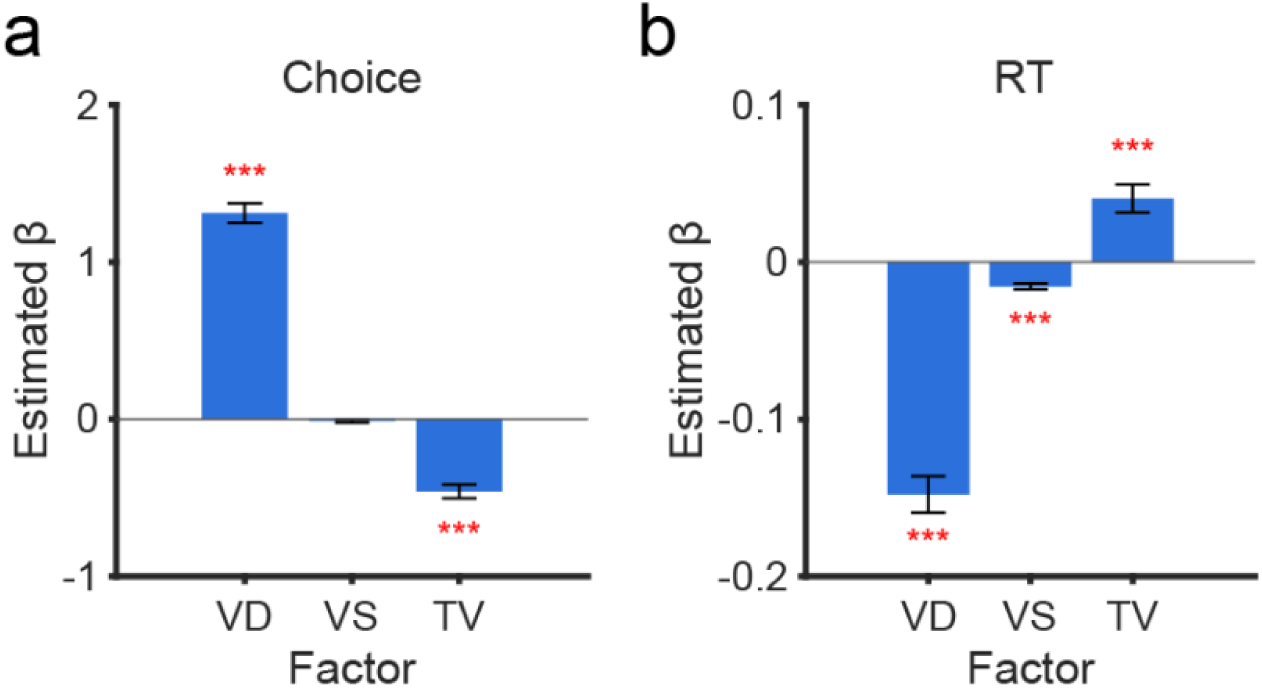
Choice accuracy and response time predicted by total WTP variability (TV). (a) Estimated fixed-effect coefficients (β ± SEM) from the GLMM predicting choice accuracy in choice task from value difference (VD), value sum (VS), and total WTP variability (TV) in value estimation task. VD positively predicted choice accuracy, whereas TV negatively predicted accuracy independently of VD and VS. ***p < 0.001. Error bars denote ±1 SEM. (b) Estimated fixed-effect coefficients (β ± SEM) from the LMM predicting response time (RT) in choice task from VD, VS, and TV in value estimation task. Larger VD and higher VS were associated with faster responses, whereas higher TV predicted longer RT. ***p < 0.001. Error bars denote ±1 SEM.

**Supplementary Figure 3.**
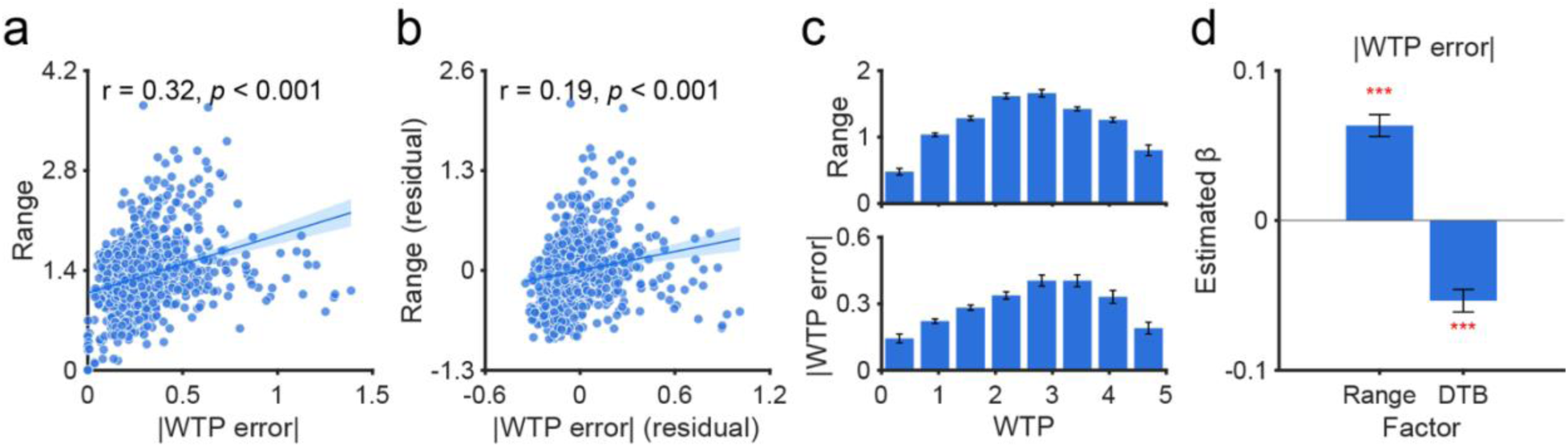
Behavioral results in the fMRI experiment. (a) Pooled scatterplot showing a significant positive correlation between WTP error and reported range across all trials and participants in the fMRI experiment (r = 0.32, p < 0.001). (b) Correlation between WTP error and reported Range after regressing out boundary distance from both variables, which remained significant (r = 0.19, p < 0.001), confirming that the positive relationship between the two variables beyond the effect of the boundary. (c) Mean reported range (top) and WTP error (bottom) as a function of WTP bin, each exhibiting an inverted-U relationship with subjective value. (d) Estimated fixed-effect coefficients (β ± SEM) from the linear mixed-effects model predicting WTP error from reported range and distance to boundary (DTB), controlling for random intercepts of participants, items, and their interaction. Reported range significantly predicted WTP error (β = 0.063 ± 0.007, p < 0.001), whereas DTB was significantly negative (β = −0.054 ± 0.008, p < 0.001), demonstrating that the relationship between range and WTP error persists after controlling for boundary proximity. *** p < 0.001. Error bars denote ±1 SEM across participants.

**Supplementary Figure 4.**
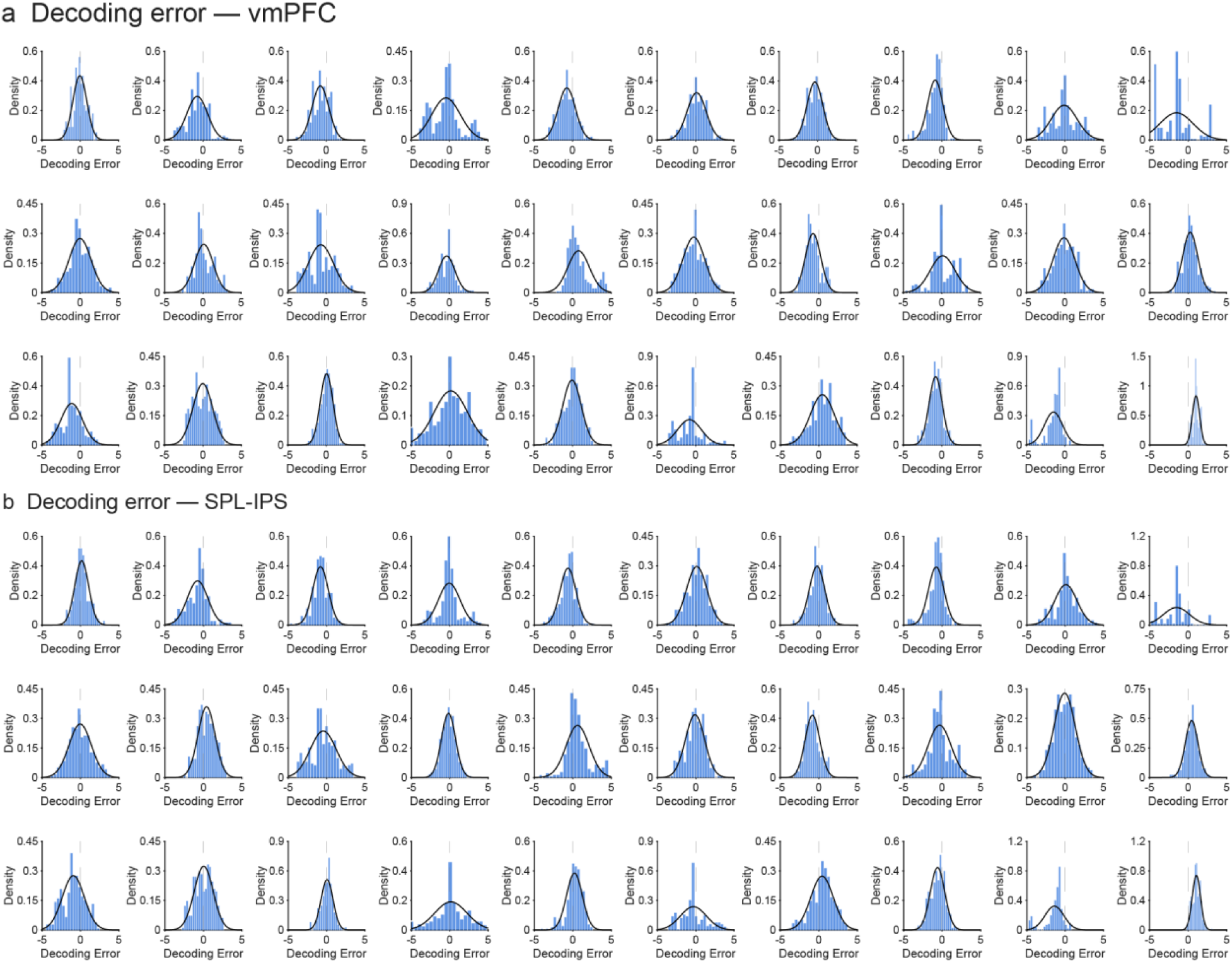
Distribution of decoding errors across participants. (a) Distribution of trial level decoding errors (decoded WTP minus item-level WTP) for each participant in vmPFC. Each panel shows the histogram and fits normal distribution for one participant. Distributions are centered near zero, indicating above chance and unbiased value decoding across participants. (b) Same as (a) but for SPL-IPS.

**Supplementary Figure 5.**
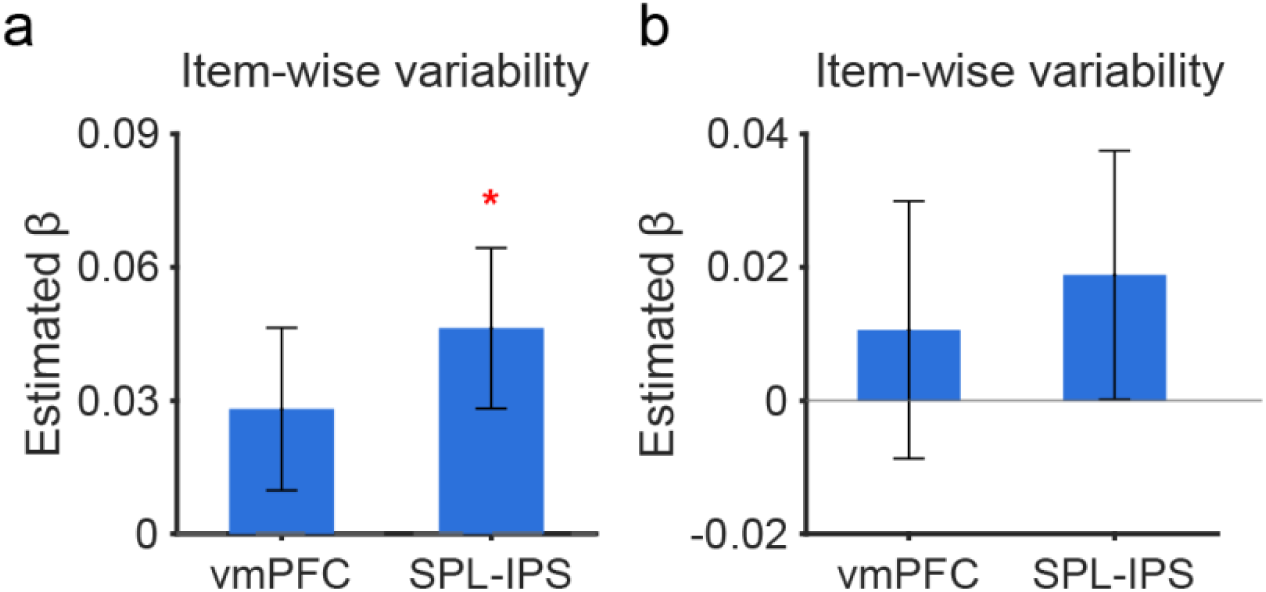
Decoded uncertainty predicts item-wise WTP variability in vmPFC and SPL-IPS. LMM-estimated β for the effect of item-wise decoded uncertainty on item-wise WTP variability across ROIs. (a) Model excluding and (b) model including item-wise boundary distance as a covariate, with random effects of participant in both models. Error bars denote ±1 SEM. *p < 0.05 after FDR correction.

## Notes

### Competing Interest Statement

The authors have declared no competing interest.

